# Neural Selectivity, Efficiency, and Dose Equivalence in Deep Brain Stimulation through Pulse Width Tuning and Segmented Electrodes

**DOI:** 10.1101/613133

**Authors:** Collin J. Anderson, Daria Nesterovich Anderson, Stefan M. Pulst, Christopher R. Butson, Alan D. Dorval

## Abstract

**Background:** Achieving deep brain stimulation (DBS) dose equivalence is challenging, especially with pulse width tuning and directional contacts. Further, the precise effects of pulse width tuning are unknown.

**Methods:** We created multicompartment neuron models for two axon diameters and used finite element modeling to determine extracellular influence from standard and segmented electrodes. We analyzed axon activation profiles and calculated volumes of tissue activated.

**Results:** Long pulse widths focus the stimulation effect on small, nearby fibers, suppressing white matter tract activation (responsible for some DBS side effects) and improving battery utilization. Directional leads enable similar benefits to a greater degree. We derive equations for equivalent activation with pulse width tuning and segmented contacts.

**Interpretations:** We find agreement with classic studies and reinterpret recent articles concluding that short pulse widths focus the stimulation effect on small, nearby fibers, decrease side effects, and improve power consumption. Our field should reconsider shortened pulse widths.

## Introduction

Rigorous efficacy testing of different deep brain stimulation (DBS) parameters requires some notion of stimulation dose equivalence. DBS programming has become more complex in recent years with the introduction of directional electrodes^1^ and interest in pulse width modulation^2–4^. Previous papers have assigned equivalence across dosages through energy, occasionally referred to as total electrical energy delivered^3,5^, charge^6^, strength-duration relationships^2^, and other quantities similar to energy^7,8^. In this work, we define equivalent doses as those consistently activating the same set of neurons. We sought to quantify the impact of pulse width tuning and directional segmentation of electrodes on the volume of tissue activated^9^ (VTA), with the goal of determining how to maintain consistent activation.

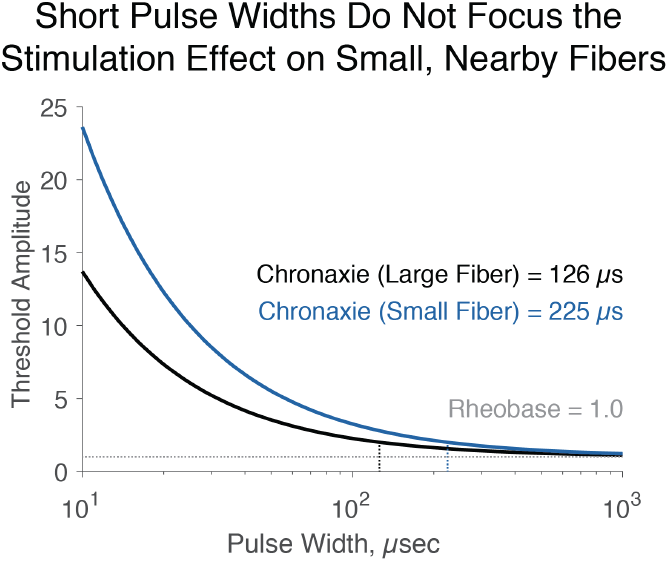
Introductory figure: Short pulse widths cannot focus the stimulation effect on small, nearby fibers. We use chronaxies from Reich et al.^2^ for large, side effect-inducing axons, and small, therapeutic axons. Strength-duration curves show that, for short pulses, threshold amplitudes diverge, and small fibers require more amplitude than large fibers, contradicting that short pulse widths could focus the stimulation effect on small, nearby fibers.

Classic biophysical studies demonstrated that short pulse widths increase selectivity for large diameter fibers^10^, and that selectivity increases with increased distance^11^. In the context of thalamic stimulation for essential tremor and subthalamic stimulation for Parkinsonism, small, nearby fibers have been associated with therapy, while distant, large fibers have been associated with side effects, supported by chronaxie values for side effects versus therapy^2^. Long pulse widths have been associated with cognitive deficits in thalamic stimulation for essential tremor^5^, while short pulse widths have been associated with increased therapeutic window^2,4,12^ and decreased side effects^3,13^. Thus, several groups have suggested that short pulse widths may focus the electrical stimulation on smaller diameter fibers near the electrode while avoiding larger fibers far away from the electrode. Not only is that suggestion at odds with the classic literature highlighted above, but it is inconsistent with strength-duration relationships. As seen in the introductory figure, short pulse widths create a divergence between the threshold amplitudes of large and small fibers, with selective activation of large, not small fibers. Interestingly, recent work suggested that pulse width tuning may be superior to directionally segmentation in side effect avoidance^3^, in contrast with reports that directionally segmented electrodes may provide a highly effective means of avoiding side effects^1,14^.

Our results lead us to conclude that pulse width tuning must be appropriately balanced through a strength-duration relationship to achieve similar dosage. We provide alternate interpretations for several results regarding pulse width tuning, suggesting that benefits from short pulse width may have arisen through improper amplitude controlling — through energy or other non-dose-equivalent metrics — and smaller volumes of neuronal activation with short pulse widths. Further, we question whether some side effects associated with longer pulse widths may be avoided by careful titration of stimulation amplitude, with a major benefit of increased battery life. In-so-doing, we find that long, not short, pulse widths may actually have the greater benefit in focusing the stimulation effect on small, nearby axons; avoiding large, distant, side effect-inducing fiber tracts, and maximizing energy efficiency. We provide equations for pulse-width tuning that enable equivalent dosing by maintaining volumes of tissue activated. Finally, we demonstrate that directional electrodes, with decreased contact surface area, can focus the effect of stimulation on small, nearby axons to an even greater extent than pulse width modulation.

## Results

We used the finite element method (in SCIRun) to solve the bioelectric field problem for monopolar stimulation from Medtronic 3387 and Abbott 6173 electrodes for 0–10 V and 0–10 mA amplitudes, respectively. We incorporated those results into the extracellular field of standard multicompartment myelinated axon models (in NEURON) projecting tangentially around the electrode, with 2.0 and 5.7 µm diameters. From these simulations, we reexamine the relationships between stimulation amplitude, pulse width, strength duration curves, contact size, energy efficiency, charge delivered, and the different volumes of tissue activated when considering 2.0 versus 5.7 µm diameter fibers.

### Long pulses and small contacts focus stimulation on small, nearby axons

We began by exploring the ability of pulse-width tuning to differentially activate axons of different sizes. First, we found maximum distances at which 2.0 and 5.7 µm axons were activated by −1.0 V, 60 µs pulses from a cylindrical electrode: 1.31 and 2.06 mm, respectively (Fig. 1A). We shortened the pulse width to 30 µs and found that the large, distant fiber was activated with less voltage than the small, nearby fiber. With a 90 µs pulse, the small, nearby fiber was activated with less voltage. Tracking axonal firing thresholds as a function of distance, we found that lengthening the pulse width progressively increases selectivity for small, nearby axons (Fig. 1B).

**Figure 1:**
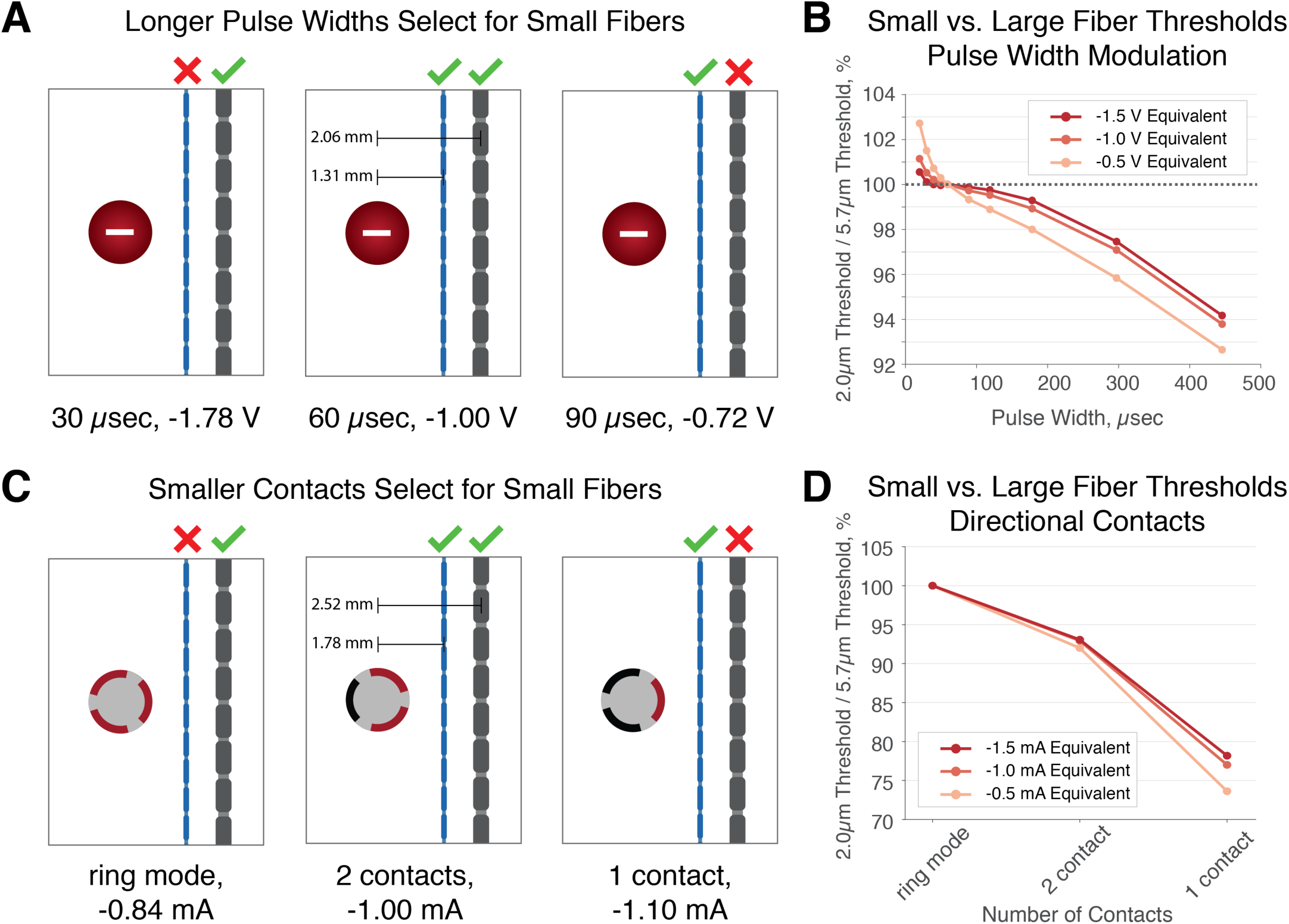
Longer pulses and smaller contacts focus the stimulation effect on small, nearby fibers. **A.** We determined maximum distances where 2.0 and 5.7 µm axons are activated by −1.00 V, 60 µs pulses from the Medtronic lead, then found minimum amplitudes to activate axons at these distances with 30 and 90 µs pulses. 30 µs stimulation more easily activated the large fiber, while the opposite was true with 90 µs. **B**. Longer pulses yield greater focusing of stimulation effect on small, nearby fibers, and the effect is greater for smaller voltages. **C**. We determined maximum distances at which 2.0 and 5.7 µm axons are activated by −1.00 mA, 60 µs pulses per contact from the Abbott lead using 2 segmented contacts, then found thresholds with ring mode and 1 contact. Ring mode more easily activated the large fiber, while one segmented contact focused the effect on the small fiber. **D**. Small fiber selectivity increases as contact surface area decreases, and the effect was greater for smaller amplitude currents. Note the different scale from B to D.

We next examined the extent to which the small-fiber selectivity achievable with pulse-width tuning could be improved with directionally selective stimulation from segmented contacts. We found maximum distances at which 2.0 and 5.7 µm axons were activated by −1.0 mA, 60 µs pulses from two segmented contacts on the same level of a directionally segmented electrode: 1.78 and 2.52 mm, respectively. Reverting to ring mode, we found that the large, distant axon was activated with less current than the small, nearby axon. With one segmented contact, the small, nearby axon was activated with less current (Fig. 1C). In plotting the threshold of the small, nearby axon relative to the large, distant axon, it is clear that decreasing the number of segmented contacts focuses the stimulation effect on small, nearby axons to a degree that is even more powerful than lengthening the pulse width (Fig. 1D, note scale change in ordinate axis from 1B).

We fit strength-duration curves to our simulation results for 5.7 and 2.0 µm diameter axons at several distances from the electrode. The shapes of the curves for different axon sizes were similar, and the relative selectivity for small axons improved with pulse width and contact proximity (Fig. 2A). We further derived strength-duration curves using one, two, and all three segmented contacts on the same level of a directionally segmented electrode. Once again, longer pulse widths and decreased separation improved selectivity for smaller axons in all contact configurations, as did reducing the number of contacts (Fig. 2B). Integrating the results for this section, selectivity for smaller fibers is best achieved by segmented contacts in close proximity to the fibers of interest, delivering long stimulation pulses.

**Figure 2:**
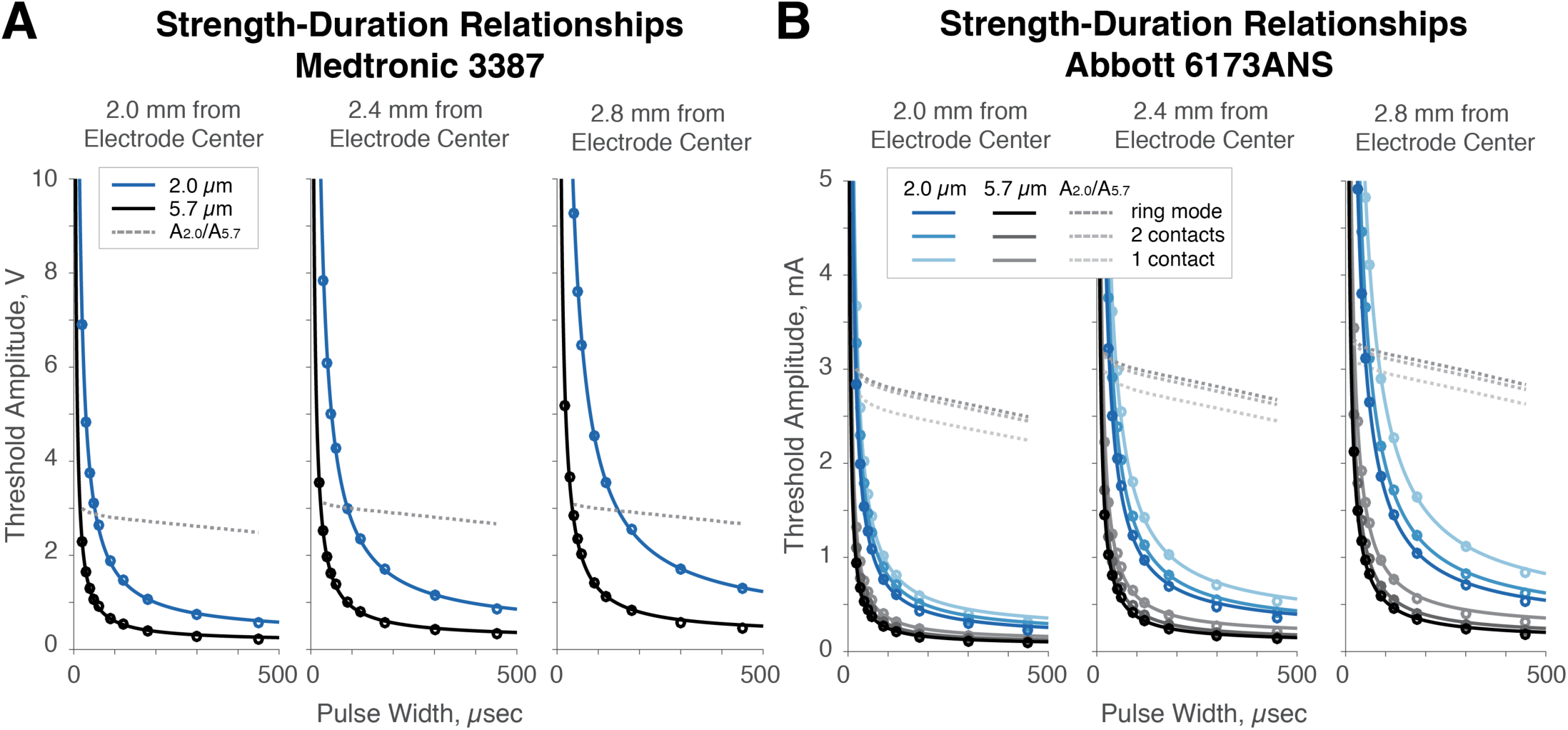
Strength-duration curves reveal that long pulse widths, proximity to electrode, and small, segmented contacts reduce preferential activation of large fibers. **A.** We derived strength-duration relationships for 5.7 and 2.0 µm fibers at 2.0, 2.4, and 2.8 mm from the electrode center. The ratio of amplitudes needed to activate 2.0 vs 5.7 µm fibers (A_2.0_/A_5.7_) decreases with distance. **B.** We derived similar relationships for 5.7 and 2.0 µm fibers at the same distances for ring mode, two, and one segmented contacts. Similar results were found as in A, and using fewer segmented contacts also improved selectivity for small versus large diameter fibers.

### Long pulses and small contacts minimize energetic demand, improving battery life

Given that longer pulses and segmented contacts improved selectivity for smaller axons, we evaluated the energy efficiency of those stimulation waveforms and configurations. When constrained to a strength-duration curve, a pulse width equivalent to the chronaxie minimizes energetic demand and maximizes theoretical battery life (Fig. 3A; see supplemental material for mathematical derivation, different from those previously published^15,16^). Specifically, given a chronaxie of ∼341 µs, shortening pulse width from 60 to 30 µs increased energetic demand by 71%, while further shortening to 20 µs increased it by 143%. Lengthening pulse width from 60 to 150 µs decreased energetic demand by 40%, while further lengthening to our chronaxie decreased it by 49%.

**Figure 3:**
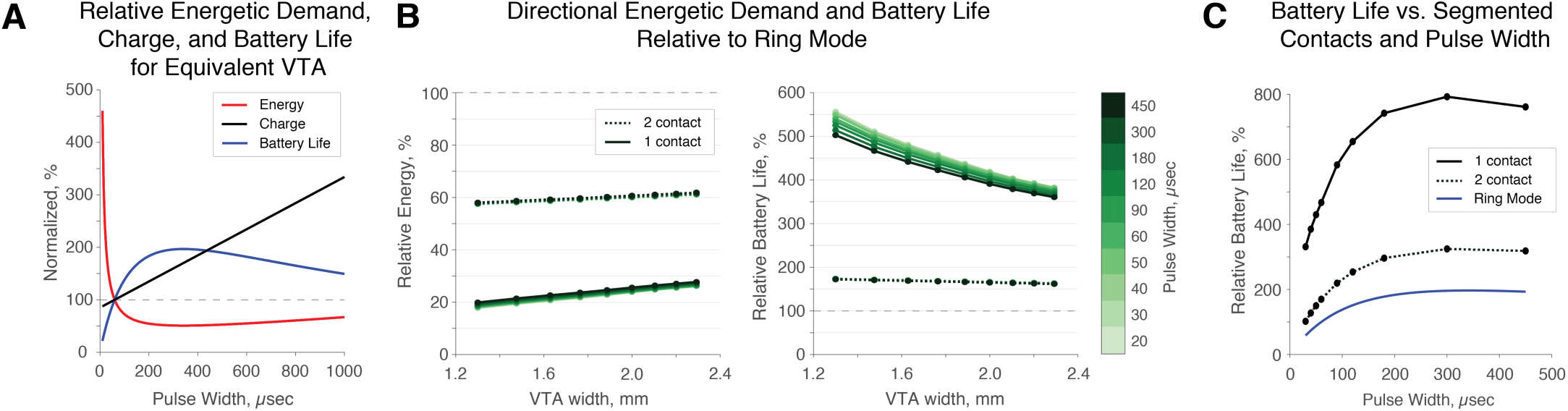
Energy efficiency and theoretical battery life are optimized by lengthening the pulse width to the value of the chronaxie and using fewer segmented contacts. **A.** Moderately long pulses (specifically, the chronaxie) optimize energy efficiency, while charge increases linearly with lengthened pulses. **B.** Energy is reduced substantially by dose-equivalent stimulation (same spread in the intended direction) with fewer segmented contacts. Theoretical battery improvements when switching to fewer contacts are greater at short pulse widths due to poor efficiency with short pulses. **C.** For a given spread in the intended direction, single contact stimulation at a moderately long pulse width maximizes battery life.

When constrained to a strength-duration curve — maintaining activation of a 2.0 µm axon oriented perpendicularly from the center of the active portion of the electrode — decreasing the number of active segmented contacts substantially decreased energetic demand (Fig. 3B, left). In general, stimulating with one contact to create an equivalent spread of activation in the intended direction decreased energy utilization by ∼75%, while using two contacts decreased it by ∼40%. This relative effect is slightly greater for very short pulses (Fig. 3B, right) for which circumferential stimulation is particularly inefficient (Fig. 3A). In absolute terms however, longer pulse widths on single contacts minimize energetic demand.

### Tuning stimulation to maintain the Volume of Tissue Activated

In this section, we seek to understand how tuning the stimulation configuration or waveform parameters will modify the activation patterns differently for neurons of different sizes. We modeled activation for 2.0 and 5.7 µm axons from a Medtronic 3387 electrode: across voltage amplitudes from −1 to -7 V for a standard 60 µs pulse width, and for a standard −2 V amplitude across pulse widths from 20 to 450 µs. Increases in voltage at a given pulse width, and increases in pulse width at a given voltage, increased stimulation spread, including the VTA and width of excitation spread from contact edge (Fig. 4A-C). Further, we modeled activation for 2.0 and 5.7 µm axons from an Abbott 6173 electrode using one, two, and three active segmented contacts: across current amplitudes from −0.5 to −3.5 mA per contact for a standard 60 µs pulse width, and for a standard −1 mA per contact across pulse widths from 20 to 450 µs. One- and two-contact cases showed greater preference for the intended direction than the opposite direction, particularly at low amplitudes and pulse widths (Fig. 4D-G). Ring-mode stimulation showed no directional preference. Importantly, modulating pulse width and modulating the amplitude via a strength-duration relationship demonstrated that pulse width modulation does not have an effect on directionality. In short, and unsurprisingly, stronger and longer stimulation increased activation, which with segmented leads, could be focused in an intended direction.

**Figure 4:**
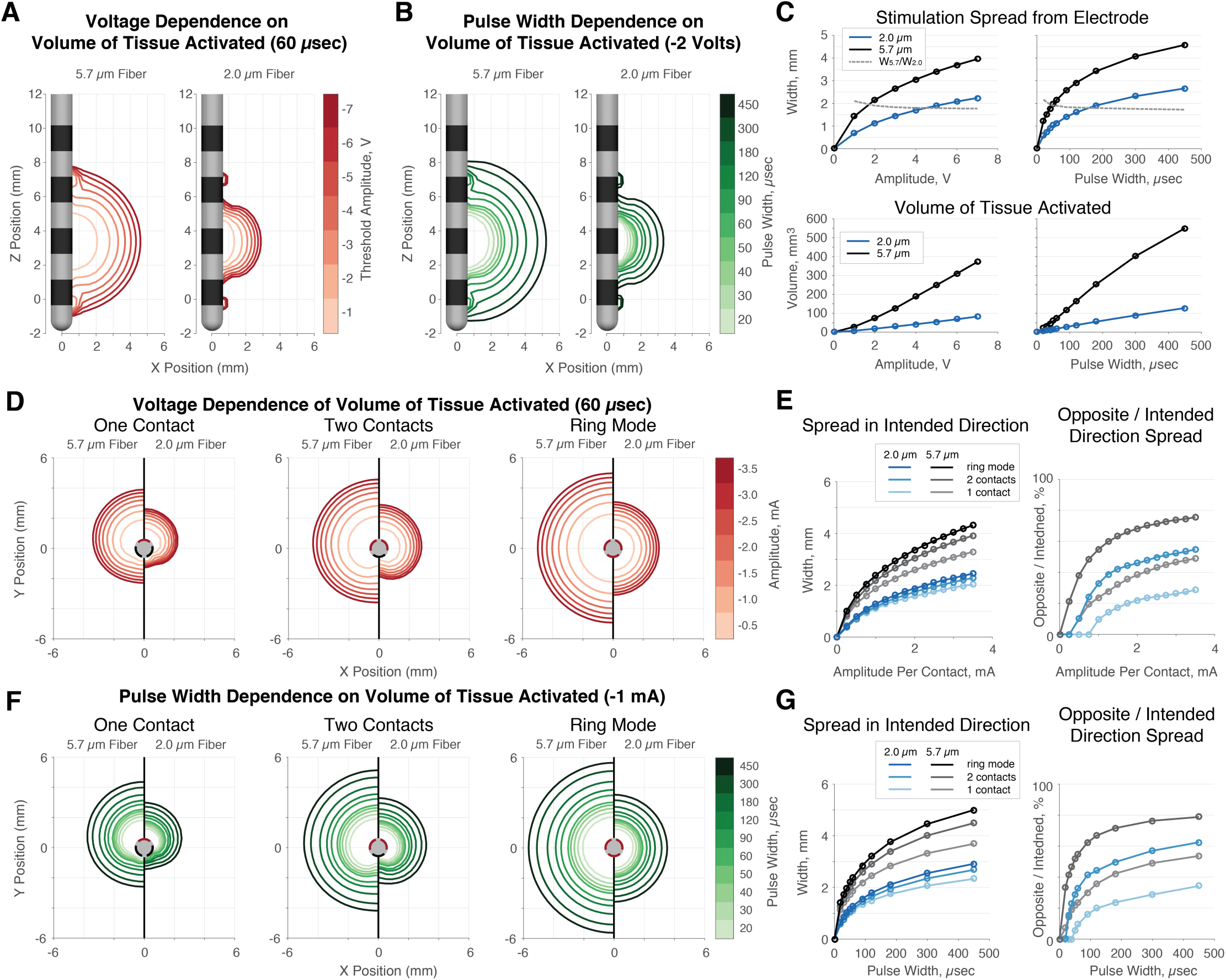
Changes in VTA with changes in voltage and pulse width. Voltage-controlled Medtronic 3387 used in A-C; Current-controlled Abbott 6673 electrode in D-G. **A.** At fixed pulse width, increasing the amplitude increases activation spread. **B.** At fixed voltage, increasing the pulse width increases activation spread **C.** At fixed pulse width, all voltages result in greater spread over 5.7 µm fibers than over 2.0 µm fibers. Dashed lines are the ratio of 5.7 to 2.0 µm fibers. Low amplitudes and pulse widths select for large fibers. **D.** Currents refer to that applied per contact. Using fewer contacts and amplitude improves directionality. **E.** Intended direction spread increases with more contacts, more pronounced for large fibers. Using fewer segmented contacts better steers stimulation in the intended direction. **F-G.** At fixed current, −1.0 mA / contact, using fewer contacts and generating a smaller VTA maximizes directionality.

Aware that we could trade pulse duration for amplitude, we determined relationships between pulse width and voltage to maintain energy, charge, and VTA size equivalence for 2.0 and 5.7 µm fibers, starting from a 60 µs pulse (Fig. 5A). At short pulse widths, energy equivalence required less voltage than VTA equivalence, which required less voltage than charge equivalence. Thus for short pulse widths, maintaining energy equivalence creates a smaller VTA, while maintaining charge equivalence creates a larger VTA. The opposite results were found for long pulse widths (Fig. 5B), supporting their improved energetic efficiency over short pulse widths when maintaining a consistent VTA.

**Figure 5:**
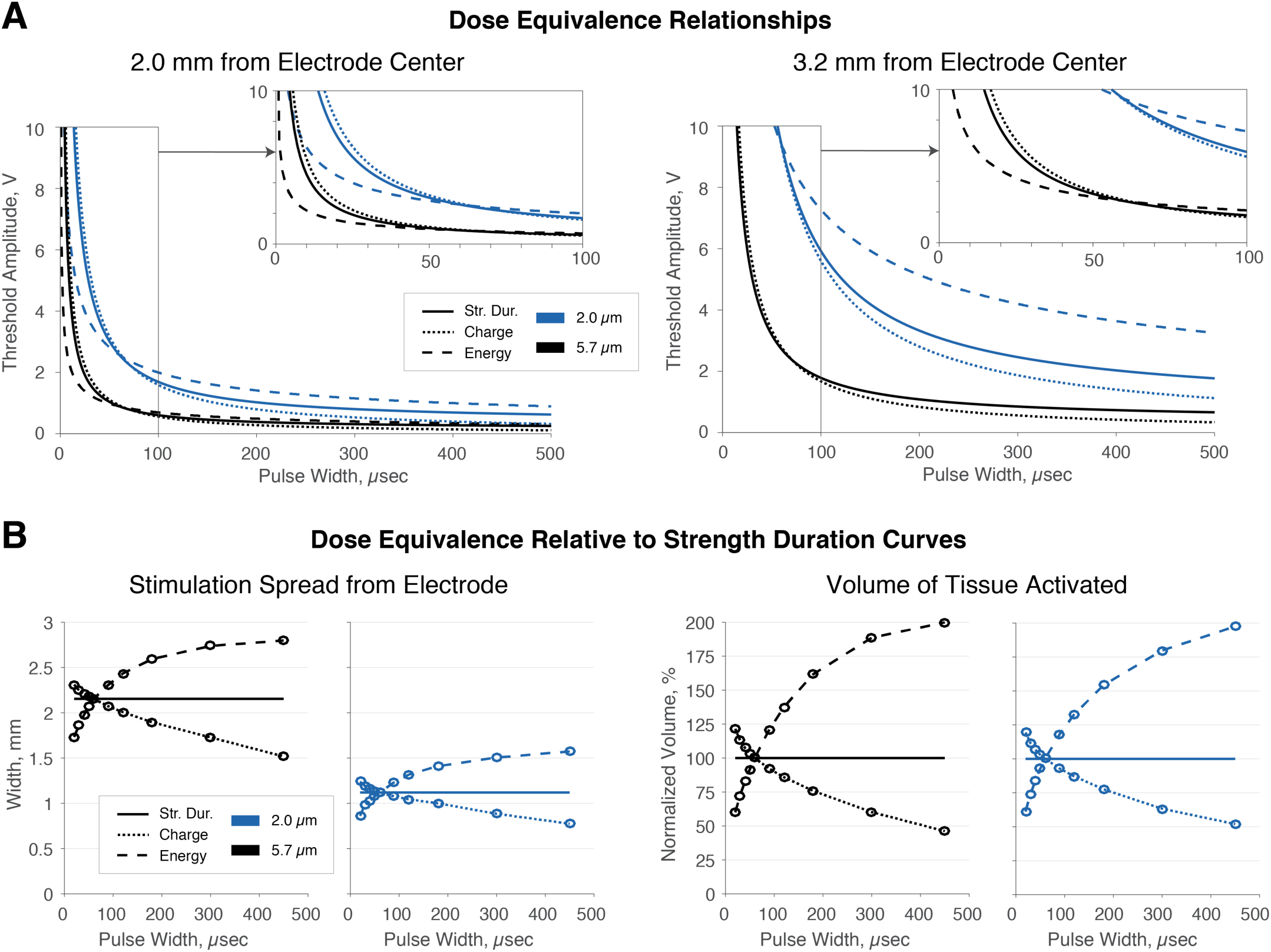
Strength-duration relationship, charge, and energy as equivalence metrics. **A.** For short pulses, the voltage required to maintain energy equivalence is less than that required to maintain a strength-duration relationship, while the voltage required to maintain equivalent charge per pulse is greater With long pulses, the trend reverses. **B.** Energy equivalence results in reduced spread of activation and VTA for short pulses and increases these for long pulses. Charge equivalence reverses this trend. Thus, both energy- and charge-equivalent parameters, with pulse width tuning, result in non-equivalent neural activation.

We sought an expression for dose equivalency that balanced amplitude and pulse width changes. Finding our 2 µm axon to have a chronaxie of 341.7 µs — well within the 200-700 µs range for grey matter^17^ — we derived the following dose-equivalency relationship (Fig. 6A): 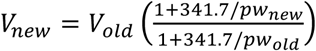. Note the dependence on chronaxie: dose-equivalent amplitudes from this relationship should be considered estimates unless modified with an appropriate chronaxie. With this relationship, voltage can be easily changed to accommodate any pulse-width tuning, while preserving the same VTA for 2 µm axons.

**Figure 6:**
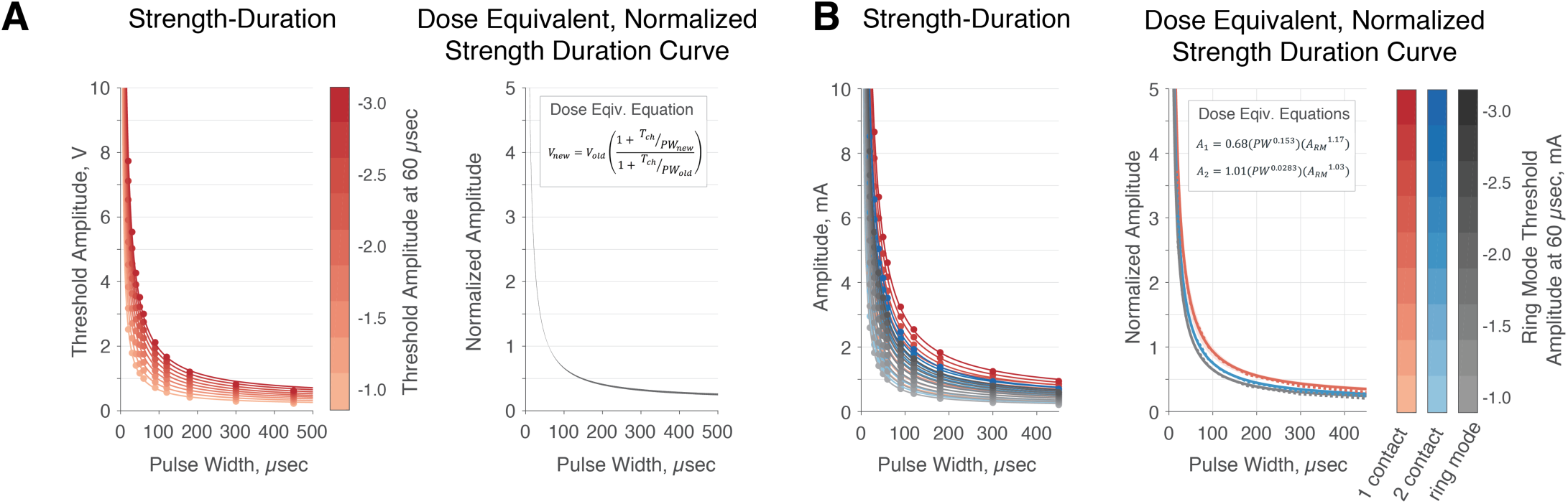
Dose Equivalence Equations and Energetic Demand. **A.** Pulse width tuning dose equivalence: we generated strength-duration curves for 2.0 µm axons at maximum distances activated by 60 µs pulses at −1.0 to −3.0 V. On right, we normalized these curves and plotted the average, the tight cloud representing standard error. **B.** Dose equivalence from ring mode to fewer contacts: for ring mode, two contacts, and one contact, we generated strength-duration curves for 2.0 µm axons and maximum distances activated by 60 µs ring mode DBS at −1.0 to −3.0 mA per contact. We predicted amplitude to activate these axons with the equivalence equations. On right, we normalized with respect to ring mode. Dashed lines indicate predictions, closely matching model data.

We next fit an equivalence equation for converting the stimulation amplitude in response to changing from ring mode to one or two segmented contacts, as in eqn. 4 (Fig. 6B): *V*_1 *contact*_ = *c* * (*pw*)*^d^*\* (*V_ring mode_*)^*b*^. In this derivation, we found the parameters b, c, and d to fit best with 1.16645, 0.68131, and 0.15298 for ring mode to one segmented contact, R^2^>0.99. For ring mode to two contacts, we found the b, c, and d as 1.02983, 1.00984, and 0.02829, R^2^>0.99 again. We used these relationships to predict equivalent currents for identical spread from ring mode to one or two segmented contacts for ten pulse widths (20–450 µs) and nine distances from the electrode, set at the maximum activation distances for 60 µs pulses of −1.0 to −3.0 mA in 0.25 mA intervals. For 20 µs, amplitudes applied in ring mode ranged from about −2.60 to −8.06 mA, while for 450 µs, the amplitudes in ring mode ranged from about −0.21 to −0.60 mA. We predicted equivalent current values when switching from ring mode to one or two contacts and compared these to model-derived currents needed to activate neurons at the given distances with the given pulse width. Across the parameter set, we found an average error in predicted equivalent current of 0.65% and a maximum of 2.52% when converting from three contacts to one. When converting to two contacts, we found an average error of 0.38% and a maximum of 2.00%. Thus, by obeying the relationship we listed above, we could reduce the number of active segmented contacts while maintaining a consistent width of neuronal activation in the intended direction.

### Analysis of Reich et al., 2015: pulse width modulation does not modify stimulation spread for side effects or rigidity control

Across both rigidity control and contraction thresholds, reported combinations of pulse width and threshold yielded insignificant changes in VTA (p=.199, rigidity control; p=.105, contractions; Pearson *t*-test) or spread (p=.129, rigidity control; p=.095 for contractions; Pearson *t*-test) (Fig. 7A). As reported, 20 µs pulses failed to generate contractions in most patients due to a 10 mA limitation. Thus, we analyzed contraction thresholds excluding 20 µs data, finding higher p-values (p=.277 for volume, p=.265 for width, Pearson *t*-test). Given that the results of the monopolar review followed the strength-duration curve of dose equivalence, pulse width modulation did not have a significant effect on VTA for either rigidity control or side effects. We conclude that pulse width modulation only increased the range of stimulation amplitude; however, the therapeutic window is unchanged (and based on our modeling results, should be slightly reduced at low pulse widths) from a biophysical perspective. The interpretation that short pulse widths focus the stimulation effect on small, nearby fibers is not supported; in fact, our data presented in the introductory figure, Fig. 1A, and Fig. 2A demonstrate the opposite.

**Figure 7:**
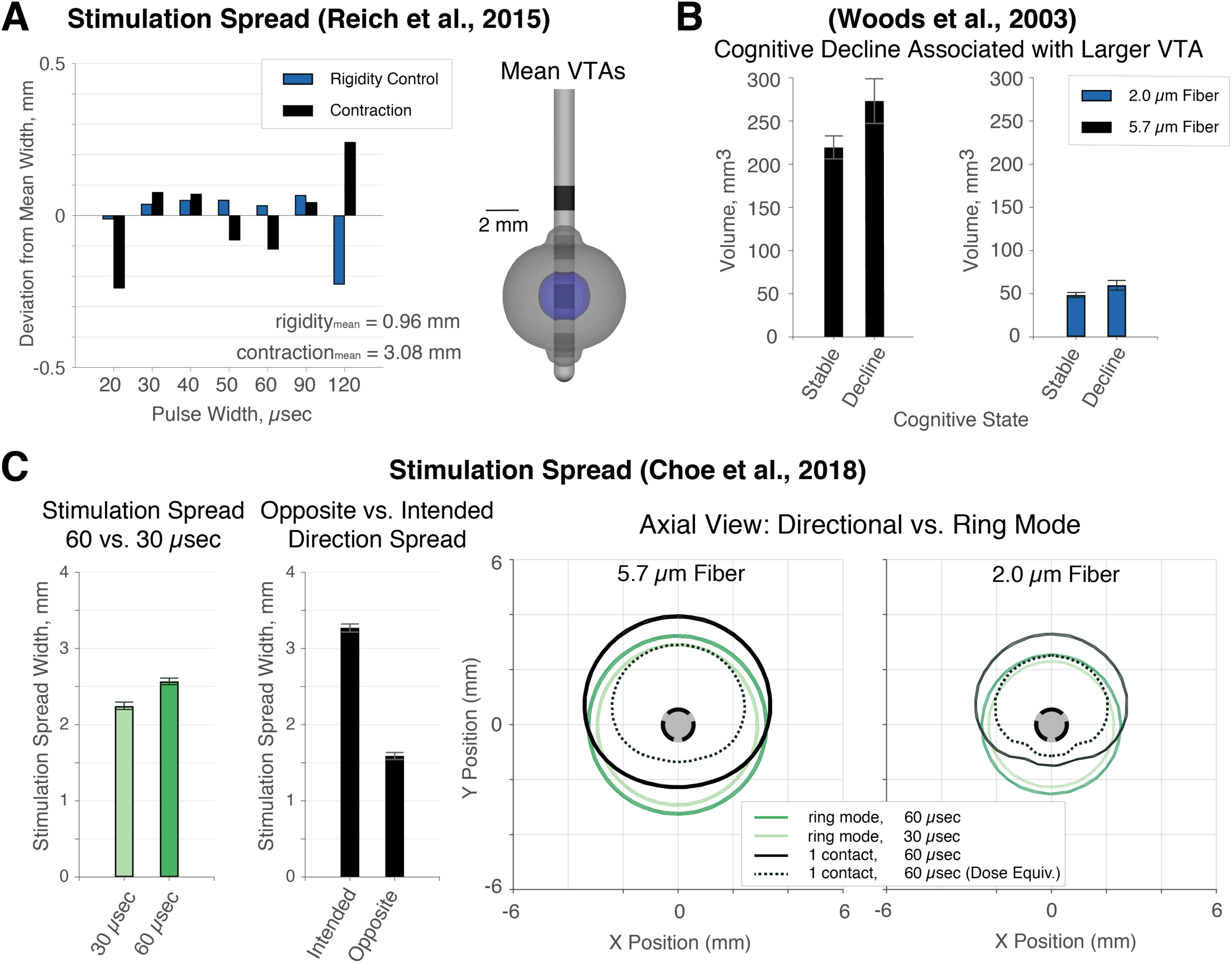
Evaluation of example and human data sets. **A.** Reich et al.^2^ report that short pulse widths increase therapeutic window and better focus the stimulation effect within small, nearby axons. In our simulations of their therapeutic and side effect thresholds, activation spread did not vary significantly across pulse widths. Pulse-width tuning therefore did not impact therapeutic window from a biophysical perspective. **B.** Woods et al.^5^ associated long pulse widths with cognitive decline in thalamic DBS. Simulations with their parameters yielded larger VTAs in those with decline than in those without, supporting that VTA size (cf. pulse width per se) may be predictive of cognitive decline. **C.** Choe et al.^3^ found that, when balanced for energy use, short pulses reduce side effects, but directional stimulation does not. From our simulations of their parameters, overall spread reduced with pulse width for both fiber sizes. Maintaining amplitude with one contact increased spread in the intended direction, but also largely in the opposite direction. Using our equivalence equation we match the intended spread for 2.0 µm fibers with more directionality and reduced 5.7 µm spread.

### Analysis of Woods et al., 2003: Patients with cognitive decline had larger VTAs

Woods et al.^5^ found that thalamic DBS patients with cognitive decline had, on average, ∼33% longer pulse width than those without. We modeled VTAs for large and small fibers with their average reported settings. VTAs were significantly smaller in those without cognitive decline than in those with (p=.027, *t*-test, Fig. 7C). We propose that another possible mechanism by which cognitive decline occurred was the larger spread of stimulation in patients with cognitive decline, potentially reaching more side effect-inducing fibers.

### Analysis of Choe et al., 2018: reduced side effects with short pulse widths may have resulted from smaller VTAs; maintaining energy equivalence with directional contacts reduces directionality

Choe et al.^3^ showed that decreasing thalamic pulse width from 60 to 30 µs while modulating amplitude to maintain energy equivalence resulted in reduced ataxic side effects. We found that activation spread was significantly greater for 60 µs parameters than for 30 µs (p=.000042, *t*-test) (Fig. 7B, left). We propose an alternate conclusion that the 30 µs case reduced ataxic side effects because the spread of activation was substantially reduced from the 60 µs case, decreasing activation of fibers responsible for side effects, while both parameter sets were greater than the minimum VTA spread required for therapy.

We analyzed further results showing that directional stimulation was ineffective for side effect avoidance. Maintaining energy equivalence from ring mode stimulation to single-contact directional stimulation results in loss of the electrode’s capability to avoid stimulation in the unintended direction (Fig. 7B, right). Axons of both sizes are activated in the intended direction across greater distances (p=.000013, *t*-test), and energy-equivalent stimulation results in ∼50% of the spread from ring mode in the opposition direction. The overall spread across intended and opposite directions is over 90% of the original linear spread, just shifted slightly in the intended direction. We propose an alternate conclusion: directional stimulation failed to generate consistent reductions in side effects because energy equivalence is particularly ineffective when switching between numbers of contacts. In the figure, we show parameters that may have been a more effective means of maintaining spread in the intended direction from the ring-mode case, generated from our segmented contact dose equivalence equation.

## Discussion

In this work, we demonstrate that short pulse widths increase selectivity for large, distant axons, while long pulse widths focus the stimulation effect on small, nearby axons, in agreement with classic literature^10,11^. Large, distant axons are generally associated with side-effects; whereas small, nearby axons are generally associated with therapeutic benefits. Further, longer pulse widths require less energy to generate given VTAs compared with shorter pulse widths. Thus, longer pulse widths preferentially activate axons responsible for therapeutic benefit while prolonging battery life. While these effects are potentially quite useful for patient programming, we demonstrate that directional segmentation of electrodes can provide these benefits to a much greater degree.

### Pulse width and therapeutic window

Short pulse widths widen the therapeutic window of DBS in terms of amplitudes. However, Reich’s data shows that the ratio between the minimum and maximum amplitudes in the therapeutic window did not significantly change over pulse widths from 20 to 120 µs^2^, and our results show that longer pulse widths can substantially change this ratio. While amplitude is the simplest way to define therapeutic window^19^, we believe that the field could be misled by continuing to strictly define the therapeutic window on one programming variable whose effects can be countered by accommodations of other programming variables (i.e. amplitude and pulse width). Instead, perhaps the field should consider therapeutic window from a biophysical perspective, based on the specific volume or set of neurons that are activated by a combination of parameters. While this is a substantially more complicated manner of thought, it will encourage understanding of the interactions of different DBS programming variables. For this more sophisticated definition, there is only a minor difference in the therapeutic windows generated with different pulse widths, with longer pulse widths enabling greater spread over small axons before reaching large axons at a set distance. While it is unclear precisely why this occurs, it could stem in part from short pulse widths requiring high voltages, which increases the second difference, based on the cable equation. Larger fibers have more surface area and reduced membrane resistance; thus, they have reduced time constants, and can could possibly respond more rapidly to a shorter stimulation pulse. Thus, for shorter pulse widths, larger diameter fibers would be even more likely to fire during the short duration.

### Longer pulse widths may have more benefits than shorter pulse widths in many DBS cases

We argue that the reduction shown in side effects with short-pulse stimulation with energy equivalence^3^ and with voltages averaging substantially lower than energy equivalence^13^ may simply arise from a non-equivalent, reduced spread of neural activation resulting in fewer side effect-inducing distant, large axons activated. However, short pulses do have some benefit. Given biophysical dose equivalence, they decrease charge injection, a nearly 10% reduction found when converting from 60 to 30 µs^2^. Very long pulse widths and interphase intervals can result in incomplete charge recovery by the charge-balancing pulse, bringing safety concerns^20,21^. However, it is unclear that shortening the pulse width below 60 µs will have much positive impact on stimulation safety, and pulse widths are sometimes lengthened to 450 µs clinically without obvious negative effects. Given that implantable pulse generators are programmed across specified increments, small pulse widths provide more steps through the therapeutic window. However, adding smaller amplitude steps to existing DBS programming technology could easily mitigate this. These advantages must be weighed against the advantages of longer pulse widths. Longer pulse widths greatly decrease energetic demand, increasing time between battery replacement surgeries. Additionally, longer pulse widths decrease preferential activation of larger fibers at distance, beneficial in many Parkinson’s and essential tremor cases.

### Long, not short, pulse widths improve battery life; battery life does not vary inversely with charge

Numerous reports have asserted that short pulse widths improve due to reduced charge. Charge scales with pulse width, but decreasing charge does not necessarily increase battery life. Battery life depends on energy, the multiple of charge and voltage, not just charge. Many batteries output constant voltage and, thus, energetic demand scales with charge; this is why many batteries can list their capacity in some variant of ampere-hours, just a multiple of coulombs. However, this requires a fixed voltage, not the case with DBS. Decreasing pulse width below standard values decreases charge, but maintaining similar neural activation requires substantially increased voltage, raising energetic demand. On the other hand, increasing pulse width increases charge but decreases voltage enough to reduce energetic demand. However, because of the rheobasic amplitude, one cannot increase pulse width indefinitely. Past the chronaxie, increasing pulse width does not result in enough voltage decrease to overcome the increased charge, and energetic demand increases. Thus, the energetically optimal pulse width is the chronaxie, 200-700 µs in grey matter^17^.

### Programming directional electrodes and gaining their benefits

In the context of directionally segmented electrodes, it is particularly important to provide dose-equivalent parameters, rather than modulating amplitude through equivalence in measures like energy. Applying the same amplitude from ring mode on only one portion of a segmented electrode can result in a large degree of stimulation around the back of the electrode, reducing the utility of the directional electrode. Achieving equivalent stimulation in the intended direction for one contact depends on creating a VTA with similar spread to that achieved in ring mode, which requires less overall amplitude. Further, we found that segmented contacts decrease selectivity for distant, large fibers. This may result from decreasing the effective width of the stimulation source, introducing a more rapidly changing voltage profile along tangential axons. Smaller diameter fibers with shorter internodal spacing between nodes are able to respond to the rapidly changing spatial field, while large diameter fibers, which can still respond more easily due to greater intermodal spacing, will be less sensitive to rapid spatial changes in electric potential, and this may effectively decrease the preference for activation of large diameter fibers.

Another benefit of directional stimulation is that using only one segmented contact decreases energy usage. When switching from ring mode to a single segmented contact, the amplitude on one contact must be increased from the amplitude used for each of the three contacts to maintain equivalent spread in the intended direction. However, the new amplitude is substantially smaller than the previous sum across three contacts. Since the amplitude term is squared in power and energy, the energy savings are substantial with one contact instead of three.

### Potential caveats

Results derived from computational models must be clinically validated. However, the models used here are based on validated models and have been well studied. Given that every patient’s lead implantation is slightly different, our dose equivalence equations should function primarily as providing initial conditions from which to determine optimal amplitude. If a chronaxie estimate is incorrect, the voltage required for an equivalent dose will vary, particularly with large pulse width changes. One must consider charge density safety when increasing pulse widths. Charge density should not exceed 30 µC/cm^2^; this has not been an issue on standard electrodes with pulse widths commonly reaching 450 µs in some indications. However, as we move towards segmented contacts and surface areas decrease, one must continue to evaluate charge densities, which rise with increased pulse width and decreased stimulation surface area.

### Conclusions

Strength duration-based dose equivalence metrics, rather than energy or charge injected, must be used to maintain equivalent neural activation and properly controlled studies with appropriate conclusions. Future work studying the effects of modulating parameters must consider how to maintain consistent VTAs. Lengthening the pulse width to match the chronaxie maximizes battery life through energy minimization and decreases preferential selection of large, distant fibers that can cause side effects, widening the therapeutic window from a biophysical standpoint. Directional stimulation through smaller, segmented contacts reduces energetic demand, effectively steers stimulation, and may also decrease the preferential selection of large, distant fibers, similar to using longer pulse widths. Finally, we provide dose equivalence equations to serve as a guide for how to modulate stimulation amplitude in response to pulse width modulation or change in number of segmented contacts. Future work will be completed to clinically test the claims made in this paper.

## Methods

### Overview

We modeled stimulation influence on activation volumes through multicompartment NEURON models, with fibers arranged tangentially around the electrodes. We determined activation profiles for 2.0 and 5.7 µm diameter neurons for the Medtronic 3387 and Abbott 6173 electrodes at pulse widths from 20 to 450 µs and amplitudes from 0 to 10 mA or 0 to 10 V, depending on whether the lead geometry is voltage or current controlled. We calculated the total charge injected and energy for each parameter set. We analyzed the relationships between pulse width and directional segmentation on the shaping of volumes of tissue activated (VTA)^22,23^ for each fiber size.

### Finite element model

We used the finite element method, implemented in SCIRun 4.7 (Scientific Computing and Imaging (SCI), Institute, University of Utah, Salt Lake City, UT), to solve the bioelectric field problem. Electrode contacts were modeled as ideal conductors, and electrode shafts were modeled as ideal insulators^24^. The volume of tissue surrounding the electrode was modeled using isotropic conductivities, using 0.2 S/m for tissue, and 0.1 S/m for the 0.5 mm encapsulation layer^25^. We solved for the electric potential solution for pulse widths of 20, 30, 40, 50, 60, 90, 120, 180, 300, and 450 for each contact on the Medtronic 3387 and Abbott 6173 leads using −1V and −0.5 mA, respectively. The outer boundary of the computational model (100 mm x 100 mm x 10 mm) was set using Dirichlet boundary conditions to stimulate the distant anode for monopolar cathodic stimulations^26^.

### NEURON Modeling

We used multicompartment axon models to quantify neuron response to extracellular stimulation. In NEURON 7.4, we used the MRG Neuron model^27^ to model 5.7 µm diameter myelinated fibers and modified parameters of the MRG model to represent 2.0 µm diameter myelinated axons^28,29^. We simulated 2.0 µm diameter fibers to quantify therapeutic activation and 5.7 µm diameter fibers to quantify side-effect activation^22^. For the Medtronic 3387 lead, tangential neurons were distributed evenly around the electrode in 0.4 mm increments from 0 to 10 mm in the radial direction and across a range of 20 mm in the axial direction. Electric potential solutions are interpolated onto each node, paranode, and internode segments of the MRG model. Each axon model is simulated over 250 ms at 130 Hz stimulation, and a binary search algorithm to determine the firing threshold with ∼0.01 V resolution. To generate a volume of tissue activated, the voltage data is rotated at 2 degree steps around the lead to take advantage of the axisymmetry of the cylindrical leads. For the directional Abbott 6173 lead, tangential neurons were arranged in 0.4 mm increments on the xy-plane through the first row of directional contacts. No axons were spaced in the axial direction to reduce computational time, and for this reason, we do not quantify volumes for the directional lead. Based on prior work by Anderson et al.^30^, cathode stimulation preferentially activates passing axons, and therefore, using tangential neurons in this modeling work is sufficient to describe the influence of extracellular cathodic stimulation.

### Selectivity of Pulse Width and Directional Modulation

To show the differential effect of pulse width stimulation on neural activation, we isolated 2.0 and 5.7 µm fibers that are activated at −1.0 V for the Medtronic 3387 lead at 60 µs. Using the same neurons, we calculated the firing threshold of those neurons at 30 µs and 90 µs to demonstrate that shorter pulse widths select for larger diameter fibers, and longer pulse widths select again larger diameter fibers. For directional stimulation, we isolated 2.0 and 5.7 µm fibers that are activated at −1.0 mA for the Abbott 6173 lead with two active contacts. We found activation thresholds for those same fibers during ring mode and single contact directional simulation to demonstration that smaller, directional contacts target increase selectivity for small fibers.

### Charge, Energetic Demand, and Battery Life Calculations

Charge per pulse (Q) is defined simply as (V × pw) / Z. We computed energy as (V^2^ × f × pw × 1 sec) / Z, where V is voltage, f is frequency, pw is pulse width, and Z is impedance^31^. We computed relative theoretical battery life as 100% / energy, normalized to that at 60 µs. We computed voltages for equivalent spread of activation for 2.0 µm fibers across pulse widths from 10-450 µs and determined energetic demand for these parameter combinations compared to that at 60 µs. For conversion from ring mode to directional contacts, we computed currents for equivalent spread in the intended direction for 2.0 µm fibers across pulse widths from 30-450 µs. Calculations were done using current, with conversion following Ohm’s law: V = I × Z, resulting in I^2^ × Z × f × pw.

### Dose equivalence with pulse width modulation

We generated models of dose equivalence across pulse widths based the on strength-duration relationship^32^, V = V_rh_ + (V_rh_ × T_ch_) / pw, with amplitude defined in terms of voltage, though the same relationship would hold for current-defined amplitudes. V refers to voltage amplitude, V_rh_ refers to the rheobase voltage, T_ch_ refers to the chronaxie, and pw refers to the pulse width. We chose to use 2.0 µm fibers for dose equivalence for three reasons. First, to maximize battery life, it is advantageous to operate at lower amplitudes within the therapeutic window, which is better represented by therapeutic, small diameter fibers rather than side effect-inducing, large fibers. Second, while our results indicate that a dose-equivalence curve based on small diameter fibers may further activate large, side effect-inducing fibers at lower pulse widths, this increase is slight and will likely not lead to side effect generation unless the stimulation is already at the ceiling of the therapeutic window. Finally, since our results present a strong argument that larger pulse widths should be clinically investigated, equating based on small fibers will maintain spread for therapeutic benefit while simultaneously slightly decreasing spread for large diameter fibers with larger pulse widths.

The rheobase and chronaxie remain consistent within each neuron given any combination of voltage and pulse width that induces firing. Thus, in order to establish a pulse width-based dose equivalent curve, we can define the ratio between voltage at a new pulse width and voltage at the old pulse width as follows in equation 1.

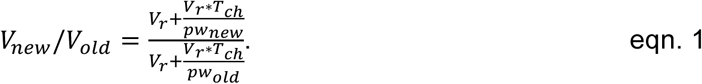

The rheobase voltage terms cancel, leading to the new voltage being defined in equation

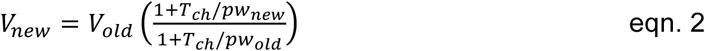

Given that different distances from the electrode define different parameterizations for the strength-duration relationship, we computed chronaxies and rheobases for 2.0 µm fibers for a number of distances with a Medtronic 3387 electrode. We found the maximum distances that enable activation of 2.0 µm fibers by 60 µs pulses at a set of voltages from −1.0 to −3.0 V, typical DBS amplitudes^33^. We found the average chronaxie for neurons in this firing window and parameterized the new voltage equation with the resulting chronaxie. Finally, we used equation 2 and a set of pulse widths from 20 to 450 µs to predict the minimum voltage required for activation at each pulse width across the set of neurons defined by being at threshold distance for activation by 60 µs pulses at −1.0 to −3.0 V. We compared the equivalence equation-predicted voltages and the modeled voltages to find the maximum error in predicted voltage required for activation across this set of fibers and apply that as confidence interval on the voltage equivalence equation.

### Dose equivalence with segmented contacts and directional stimulation

We found the maximum distance at which 2.0 µm diameter fibers could be activated by 60 µs pulses in ring mode at the lower level of segmented contacts at −1.0 to −3.0 mA applied to each contact. We then determined the minimum voltage required to activate neurons in the intended direction of stimulation at these distances using only one or two of the segmented contacts with 60 µs pulses. We used a power law to fit the relationship between voltage required with fewer contacts to voltage required by three contacts, shown in equation 3.

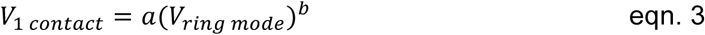

We repeated this process for 2.0 µm fibers at the same distances as above, but with pulse widths from 30 to 450 µs – there were not enough firing data points for 20 µs to fit a curve. Since the exponential term, *b*, varied quite minimally across pulse widths, and we were able to find a constant value for *b*. The coefficient, *a*, followed a power law relationship with change in pulse width, and we parameterized the coefficient *a* accordingly. Therefore, we made a final fit of dose equivalence for differing numbers of segmented contacts following the following relationship in equation 4.

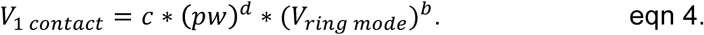

To address this possible error introduced by parameterizing *b* as the average value across different pulse widths, we used the equivalence relationship to predict spread from the electrode in the intended direction of stimulation and compared it to VTA models. We report the maximum of this error as a confidence interval on the voltage equivalence equation.

### Clinical Evaluation of dose equivalence

For previous clinical experiments that have used pulse width as an independent variable in human subjects research, we ran VTAs given reported stimulation parameters to determine whether dose equivalence was controlled for across conditions:

#### Reich et al., 2015

We analyzed the clinical data from the 2015 paper by Reich et al.^2^. We computed VTAs at thresholds for rigidity control (2.0 µm fibers) and contraction side effects (5.7 µm fibers) for all pulse widths: 20, 30, 40, 50, 60, 90, and 120 µs. For simplicity, we modeled with Medtronic 3387 electrodes, and converted current to voltage via Ohm’s law with a typical impedance of 1 kΩ. We compared VTA sizes across all combinations of pulse width and amplitudes used in Reich et al to verify dose equivalence.

#### Choe et al., 2018

We also analyzed the pulse width and directional experiment from Choe et al.. We modeled spread over 5.7 µm fibers with Abbott 6173 leads at given parameters for ring mode (−3.44 mA at 60 µs vs. −4.85 mA at 30 µs). Additionally, we found the volume of tissue activated for energy-equivalent stimulation from ring mode stimulation to single-contact directional stimulation. Additionally, we recomputed the VTA spread using strength-duration dose equivalence described in the earlier section “*Dose equivalence with segmented contacts and directional stimulation.*“

#### Woods et al., 2003

To simulate VTAs derived from Woods et al., we modeled VTAs for large and small fibers with their average reported settings at 105 µs pulse with and 80 µs pulse width (105.37 µs mean pulse width showing cognitive decline versus 79.09 µs without). We isosurfaced VTAs at the reported −3.18 V and - 3.51 V for both large and small diameter VTAs to generalize VTA spread in Woods et al., to determine the influence of activation volume on clinical outcomes.

### Statistics

When relevant, we present results as mean +/-standard error. We use the following abbreviations: Student *t*-test is referred to as *t*-test and Pearson correlation coefficient *t*-test is referred to as Pearson *t*-test.

## Supplemental Material: derivation of chronaxie as the pulse width that minimizes energy consumption for a given set of neuronal activation

We wish to maximize battery life for equivalent neural activation through modulation of pulse width and voltage. Solving how to do this is the same as asking how one minimizes energy when constrained by a strength-duration relationship, which is as follows: 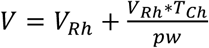, where V is voltage, V_Rh_ is rheobase voltage (the minimum voltage to activate a neuron at an infinite duration of stimulation), T_Ch_ is chronaxie (the stimulation duration required to activate a neuron at double the rheobase voltage), and pw is pulse width. Solving for pw gives 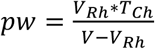.

Moving onto power and energy, *E* = *P* ∫ *dt* and *P* = *I* * *V*, where E is energy, P is power, t is time, I is current, and V is voltage. From Ohm’s Law, *V* = *I* * *R*, and, thus, *P* = *V*^2^/*R*. However, we do not constantly apply this stimulation, only at pulses for a given frequency, f. Therefore, we are only applying the stimulation at *pw* * *f* per second. Therefore we solve for energy, *E* = (*V*^2^ * *f* * *pw* * 1 *sec*)/*R*, for unit time.

From the above, constraining voltage and pulse width to a strength-duration relationship, 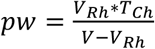. Therefore, we can substitute this in for pulse width and arrive at 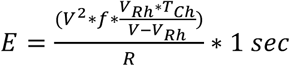 Given that we are not modulating frequency, resistance is not affected by modulating voltage, and 1 second is a constant, we arrive at normalized energy, 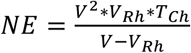. This normalized energy will scale with energy, and, thus, minimizing NE is equivalent to minimizing energy. To minimize NE in terms of voltage, we must take the derivative of NE with respect to voltage and find where this is equal to zero. Via the quotient rule, 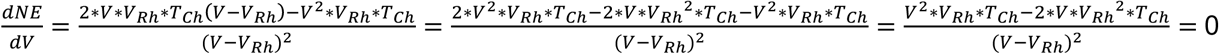. It follows that *V* * *V*_*Rh*_ * *V*_*Ch*_ — 2 * *V*_*Rh*_^2^ * *T_Ch_* = 0, constrained by *V* ≠ *V*_*Rh*_.

Therefore, either V = 0, which will not yield neural activation, or *V* * *V*_*Rh*_ * *V*_*Ch*_ — 2 * *V*_*Rh*_^2^ * *T_Ch_* = 0. For the second option, *V* * *V*_*Rh*_ * *T*_*Ch*_ = 2 * *V*_*Rh*_^2^ * *T_Ch_*. The chronaxie values cancel, and isolating V gives the following: *V* = 2 * *V*_*Rh*_. Following the strength duration relationship, if we stimulate at double the voltage, we are applying a pulse width equivalent to the chronaxie. A quick check of the surrounding energy values indicates that this indeed a minimum. Therefore, to minimize energy for a given neuronal activation, one should stimulate at the chronaxie value for pulse width and double the rheobase threshold.

## References

1. Martens, H. C. F. et al. Spatial steering of deep brain stimulation volumes using a novel lead design. Clin. Neurophysiol. 122, 558–566 (2011).

2. Reich, M. M. et al. Short pulse width widens the therapeutic window of subthalamic neurostimulation. Ann. Clin. Transl. Neurol. 2, 427–432 (2015).

3. Choe, C.-U. et al. Thalamic short pulse stimulation diminishes adverse effects in essential tremor patients. Neurology 91, e704–e713 (2018).

4. Bouthour, W. et al. Short pulse width in subthalamic stimulation in Parkinson’s disease: a randomized, double-blind study. Mov. Disord. 33, 169–173 (2018).

5. Woods, S. P., Fields, J. A., Lyons, K. E., Pahwa, R. & Tröster, A. I. Pulse width is associated with cognitive decline after thalamic stimulation for essential tremor. Parkinsonism Relat. Disord. 9, 295–300 (2003).

6. McDermott, H. & McKay, C. Comment on: Short pulse width widens the therapeutic window of subthalamic neurostimulation. Ann. Clin. Transl. Neurol. 2, 984–985 (2015).

7. Moro, E. et al. The impact on Parkinson’s disease of electrical parameter settings in STN stimulation. Neurology 59, 706–713 (2002).

8. Voges, J. et al. Bilateral high-frequency stimulation in the subthalamic nucleus for the treatment of Parkinson disease: correlation of therapeutic effect with anatomical electrode position. J. Neurosurg. 96, 269–279 (2002).

9. Butson, C. R. & McIntyre, C. C. Role of electrode design on the volume of tissue activated during deep brain stimulation. J. Neural Eng. 3, 1–8 (2006).

10. Gorman, P. H. & Mortimer, J. T. The Effect of Stimulus Parameters on the Recruitment Characteristics of Direct Nerve Stimulation. IEEE Trans. Biomed. Eng. BME-30, 407–414 (1983).

11. Grill, W. M. & Mortimer, J. T. The effect of stimulus pulse duration on selectivity of neural stimulation. IEEE Trans. Biomed. Eng. 43, 161–166 (1996).

12. Rizzone, M. et al. Deep brain stimulation of the subthalamic nucleus in Parkinson’s disease: effects of variation in stimulation parameters. J. Neurol. Neurosurg. Psychiatry 71, 215–219 (2001).

13. Kroneberg, D., Ewert, S., Meyer, A.-C. & Kühn, A. A. Shorter pulse width reduces gait disturbances following deep brain stimulation for essential tremor. J. Neurol. Neurosurg. Psychiatry (2019). doi:10.1136/jnnp-2018-319427

14. Keane, M., Deyo, S., Abosch, A., Bajwa, J. A. & Johnson, M. D. Improved spatial targeting with directionally segmented deep brain stimulation leads for treating essential tremor. J. Neural Eng. 9, 046005 (2012).

15. Grill, W. M. Model-Based Analysis and Design of Waveforms for Efficient Neural Stimulation. Prog. Brain Res. 222, 147–162 (2015).

16. Kroll, M. W. A Minimal Model of the Monophasic Defibrillation Pulse. Pacing Clin. Electrophysiol. 16, 769–777 (1993).

17. Ranck, J. B. Which elements are excited in electrical stimulation of mammalian central nervous system: A review. Brain Res. 98, 417–440 (1975).

18. Ramasubbu, R., Lang, S. & Kiss, Z. H. T. Dosing of Electrical Parameters in Deep Brain Stimulation (DBS) for Intractable Depression: A Review of Clinical Studies. Front. Psychiatry 9, (2018).

19. Reich, M. M. & Volkmann, J. Reply to comment on: Short pulse width widens the therapeutic window of subthalamic neurostimulation. Ann. Clin. Transl. Neurol. 2, 986 (2015).

20. Lempka, S. F., Howell, B., Gunalan, K., Machado, A. G. & McIntyre, C. C. Characterization of the stimulus waveforms generated by implantable pulse generators for deep brain stimulation. Clin. Neurophysiol. 129, 731–742 (2018).

21. Merrill, D. R., Bikson, M. & Jefferys, J. G. R. Electrical stimulation of excitable tissue: design of efficacious and safe protocols. J. Neurosci. Methods 141, 171–198 (2005).

22. Butson, C. R., Cooper, S. E., Henderson, J. M. & McIntyre, C. C. Patient-specific analysis of the volume of tissue activated during deep brain stimulation. NeuroImage 34, 661–670 (2007).

23. Butson, C. R. & McIntyre, C. C. Tissue and electrode capacitance reduce neural activation volumes during deep brain stimulation. Clin. Neurophysiol. 116, 2490–2500 (2005).

24. Vorwerk, J., Brock, A. A., Anderson, D. N., Rolston, J. D. & Butson, C. R. A retrospective evaluation of automated optimization of deep brain stimulation parameters. bioRxiv 393900 (2018). doi:10.1101/393900

25. Butson, C. R., Maks, C. B. & McIntyre, C. C. Sources and effects of electrode impedance during deep brain stimulation. Clin. Neurophysiol. 117, 447–454 (2006).

26. Anderson, D. N., Osting, B., Vorwerk, J., Dorval, A. D. & Butson, C. R. Optimized programming algorithm for cylindrical and directional deep brain stimulation electrodes. J. Neural Eng. 15, 026005 (2018).

27. McIntyre, C. C., Richardson, A. G. & Grill, W. M. Modeling the excitability of mammalian nerve fibers: influence of afterpotentials on the recovery cycle. J. Neurophysiol. 87, 995–1006 (2002).

28. McIntyre, C. C., Grill, W. M., Sherman, D. L. & Thakor, N. V. Cellular effects of deep brain stimulation: model-based analysis of activation and inhibition. J. Neurophysiol. 91, 1457–1469 (2004).

29. Pelot, N. A., Behrend, C. E. & Grill, W. M. Modeling the response of small myelinated axons in a compound nerve to kilohertz frequency signals. J. Neural Eng. 14, 046022 (2017).

30. Anderson, D. N., Duffley, G., Vorwerk, J., Dorval, A. (Chuck) & Butson, C. R. Anodic Stimulation Misunderstood: Preferential Activation of Fiber Orientations with Anodic Waveforms in Deep Brain Stimulation. J. Neural Eng. (2018). doi:10.1088/1741-2552/aae590

31. Koss, A. M., Alterman, R. L., Tagliati, M. & Shils, J. L. Calculating total electrical energy delivered by deep brain stimulation systems. Ann. Neurol. 58, 168–168 (2005).

32. Lapicque, L. Recherches quantitatives sur l’excitation electrique des nerfs traitee comme une polarization. J. Physiol. Pathol. Gen. 9, 620–635 (1907).

33. Kuncel, A. M. & Grill, W. M. Selection of stimulus parameters for deep brain stimulation. Clin. Neurophysiol. Off. J. Int. Fed. Clin. Neurophysiol. 115, 2431–2441 (2004).

